# Sex-linked markers in an Australian frog *Platyplectrum ornatum* with a small genome and homomorphic sex chromosomes

**DOI:** 10.1101/2022.08.09.503425

**Authors:** Chad Schimeck, Foyez Shams, Ikuo Miura, Simon Clulow, Zuzana Majtanova, Janine Deakin, Tariq Ezaz

## Abstract

Amphibians have highly diverse sex-determining modes leading to a notable interest in vertebrate sex determination and sex chromosome evolution. The identification of sex-determining systems in amphibians, however, is often difficult as a vast majority consist of homomorphic sex chromosomes making them hard to distinguish. In this study, we used Diversity Array Technology sequencing (DArTseq™) to identify the sex-determining system in the ornate burrowing frog from Australia, *Platyplectrum ornatum*. We applied DArTseq™ to 44 individuals, 19 males and 25 females, collected from two locations to develop sex-linked markers. Unexpectedly, these 44 individuals were classified into two distinct population clusters based on our SNP analyses, 36 individuals in cluster-1, and 8 individuals in cluster-2. We then performed sex-linkage analyses separately in each cluster. We identified 35 sex-linked markers from cluster-1, which were all associated with maleness. Therefore, *P. ornatum* cluster-1 is utilising a male heterogametic (XX/XY) sex-determining system. On the other hand, we identified 210 sex-linked markers from cluster-2, of which 89 were male specific, i.e., identifying XX/XY sex determining system and 111 were female specific, i.e., identifying ZZ/ZW sex determining system, suggesting existence of either male or female heterogametic sex determining system in cluster-2. We also performed cytogenetic analyses in 1 male and 1 female from cluster-1; however, we did not detect any visible differentiation between the X and Y sex chromosomes. We also mapped sex-linked markers from the two clusters against the *P. ornatum* genome and our comparative analysis indicated that the sex chromosomes in both clusters shared homologies to chromosome 10 (autosome) of *Rana temporaria* and ZWY sex chromosome of *Xenopus tropicalis*. It is plausible that the cluster-2 has a potential to be either male or female heterogamety in sex determination, requiring further investigation.

## INTRODUCTION

Vertebrate sex determination has become a fundamental area for better understanding the evolutionary advantages and disadvantages of a species. Amphibians play a crucial role in filling the knowledge gaps as they typically contain alternative sex-determination systems, even amongst geographic populations within single species (Miura, 2017; Nishioka, Hanada, Miura, & Ryuzaki, 1994; Nishioka, Miura, & Saitoh, 1993; Rodrigues, Merilä, Patrelle, & Perrin, 2014; Toups, Rodrigues, Perrin, & Kirkpatrick, 2019). The most well researched amphibian order is Anura consisting of frogs and toads (Ma & Veltsos, 2021). By understanding the sex of anurans, we can better identify the evolutionary advantages of each sex-determining system not just in frogs and toads but in all vertebrates.

The Anura comprises over seven thousand described species, representing frog and toads, distributed in tropical and temperate regions of the world (Vitt & Caldwell, 2013). Anurans are considered a model group for studying sex chromosome evolution given the presence of diverse modes of sex determination, homomorphic and heteromorphic sex chromosomes, multiple sex chromosome systems, rapid rate of turnover and sex reversal in natural environments (Jeffries et al., 2018; Ma & Veltsos, 2021; Miura, 2017; Miura et al., 2021; Nishioka et al., 1994; Ruiz-García, Roco, & Bullejos, 2021; Xu et al., 2022). Although mode of sex determination in a large number of species is yet to be discovered, a recent study reviewed sex determination in 222 anuran species and reported that the majority with known sex determination systems are either male heterogametic (XX/XY) or female heterogametic sex chromosomes (ZZ/ZW) (Ma & Veltsos, 2021). It also reported the abundance of homomorphic sex chromosomes amongst the Anura, with no cytogenetically distinguishable characteristics such as size polymorphism or heterochromatinization. Furthermore, transitions between male (XX/XY) and female (ZZ/ZW) heterogametic sex chromosome systems have also been reported in this group (such as Japanese wrinkled frog, *Glandirana rugosa*), where both XX/XY and ZZ/ZW sex chromosomes were found in different geographic populations of the same species (Miura, 2007; Nishioka et al., 1994; Ogata, Hasegawa, Ohtani, Mineyama, & Miura, 2008).

In vertebrates, sex determination is either governed by genetics (genetic sex determination, GSD) where so-called master sex-determining genes on sex chromosomes are responsible for maleness or femaleness, or by environmental factors (environmental sex determination, ESD) such as temperature (Bachtrog et al., 2014; Capel, 2017). Unlike reptiles, mammals, birds and fishes, all amphibians reported to date show solely genetic sex determination (Ezaz, Stiglec, Veyrunes, & Graves, 2006; Saidapur, Gramapurohit, & Shanbhag, 2001; Sarre, Ezaz, & Georges, 2011). To date, eight genes have been proposed as candidate sex-determining genes in frogs including *Amh, Ar, Cyp19a1, Cyp17, dmrt1, Foxl2, Sf1* and *Sox3* (Miura, 2017). However, among frogs, *Dm-W* is the only confirmed sex-determining gene found in the W chromosome of the African clawed frog *Xenopus laevis*, functioning primarily to determine ovary (Yoshimoto et al., 2010; Yoshimoto et al., 2008).

Australia is home to around 248 described frog species, which form highly diverse lineages that have adapted to and evolved across this largely arid and huge continent (Clulow & Swan, 2018). However, very little is known about sex determination in Australian frogs. Prior to the current study, only two Australian frog species have been investigated to understand their sex-determining system and identify sex chromosomes involved (Mahony, 1991; Sopniewski, Shams, Scheele, Kefford, & Ezaz, 2019). In our previous study Sopniewski et al. (2019), we identified sex linked markers in a threatened species *Litoria aurea* and discussed its implications for the benefit of sex-linked markers, not only in understanding evolution of sex determination, but also how these markers can assist in conservation and management of vulnerable species in establishing captive breeding programs.

In this study, we investigated another Australian native species, the ornate burrowing frog *Platyplectrum ornatum*, found throughout northern and eastern regions of Australia. It is of particular interest because it has one of the smallest genomes among amphibians globally (Lamichhaney et al., 2021). We applied genotyping by sequencing and cytogenetic analyses to identify sex-linked markers and sex chromosomes in this species. Our study identified two distinct population clusters within our samples, one containing a XX/XY sex chromosome system, while the other contained both XY and ZW sex chromosome systems, the first such system reported for an Australian frog. We discuss the evolutionary origins of *P. ornatum* sex chromosomes by comparative analysis of the sex-linked markers.

## RESULTS

Sequencing using Diversity Arrays Technology (DArTseq) yielded 26,549 SNPs and 33,444 PA loci in 44 individuals. The average call ratio of the SNPs loci was 0.86, and the PA loci was 0.97, indicating that all the loci were sequenced successfully in almost all individuals. The average sequencing reproducibility of the SNP loci was 0.98 and the SilicoDArT loci was 0.99. The high call ratio and reproducibility of the sequenced loci indicate the good quality of these markers for further analysis.

### Genetic diversity and population genetic structure

Sex chromosomes or the mode of sex determination can vary down to population level in frogs. Prior to performing the sex-linked marker analyses to understand sex-determination in *P. ornatum*, we performed population genetic analyses to infer whether all 44 individuals used in this study belong to a single species. We used 4,691 autosomal SNP loci (excluding all sex-linked loci) with 0% null allele and 100% reproducibility for the population genetic analyses. The STRUCTURE and Principal Component Analyses (PCA) indicated a distinct clustering of two genetic groups where 36 individuals (Cluster-1) were genetically distinct from the other eight (8) individuals (Cluster-2) (Fig 1C, E). The pairwise F_st_ value between the two clusters was 0.24. The Nei (1972) genetic distance between the two clusters was 0.13 while the mean genetic distance among individuals in Cluster-1 was 0.19 and that in Cluster-2 was 0.07.

**Figure 1:**
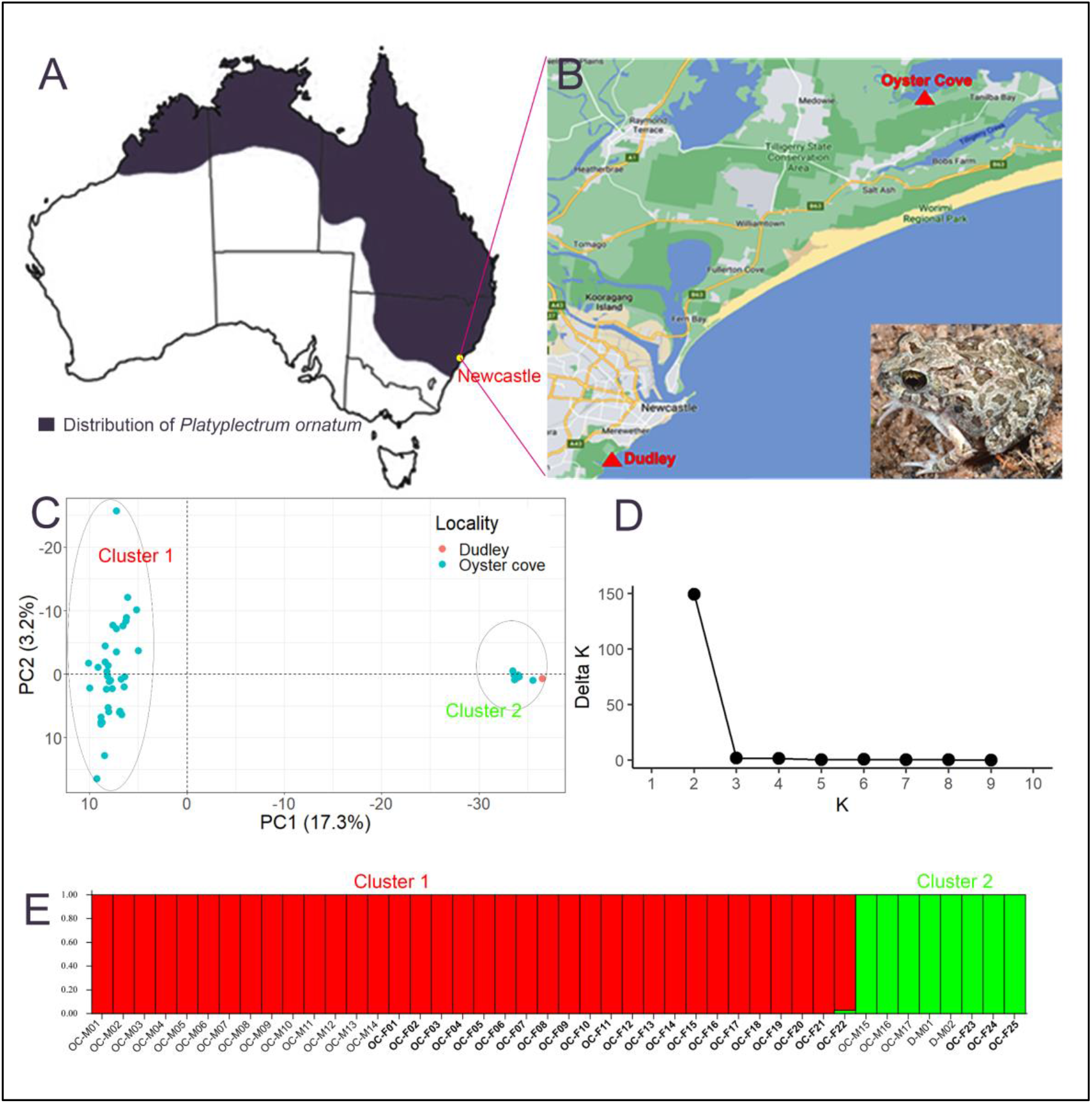
Population genetic analyses of *Platyplectrum ornatum*. A – Distribution range of the species. B – Sampling location of the individuals used in this study (in the inset, a photograph of one specimen). C – Principal Component Analysis (PCA) plot using only autosomal markers suggesting two distinct genetic groups withing the samples. D – Delta K values from the structure analysis. E – Structure plot separating two clusters within the samples. OC-M = Oyster Cove Male, OC-F = Oyster Cove Female, D-M = Dudley Male.

The Analysis of Molecular Variance (AMOVA) indicated that around 23.47% variation was described by the molecular comparison between the two clusters. The significance test of the molecular variance indicated significant molecular genetic variation between the two clusters (Table 1). This suggests that the specimens collected from Oyster cove and Dudley include a cryptic new species that inhabits sympatrically in Oyster cove. Thus, although this finding is very important for understanding speciation in Australian native frogs, we separated the two clusters and analysed those data separately to identify sex-linked markers.

**Table 1:** Analysis of Molecular Variance (AMOVA) among samples used in this study. df: degrees of freedom. *: significant

|  | df | Sum.Sq | Mean.S<br>q | Sigma | Molecular<br>variance<br>(%) | Pvalue |
| --- | --- | --- | --- | --- | --- | --- |
| Between Clusters | 1 | 11557.79 | 11557.79 | 390.84 | 23.47* | 2e<sup>-5</sup> |
| Between samples Within<br>Clusters | 42 | 55642.07 | 1324.81 | 50.56 | 3.04 | 0.11 |
| Within samples | 44 | 53842.00 | 1223.68 | 1223.68 | 73.49* | 1e<sup>-6</sup> |
| Total | 87 | 121041.86 | 1391.29 | 1665.09 | 100.00 |  |

### Sex-linked markers in *P. ornatum*

To infer the mode of sex determination, we tested all SNPs and PA loci for both male heterogametic (XX/XY) and female heterogametic (ZZ/ZW) sex determination systems criteria (see methods). In Cluster-1, the filtering of loci for sex linkage resulted in 11 SNPs and 24 PA loci (Table S1), all of which showed association for a male heterogametic sex determining system (XX/XY) in this cluster of *P. ornatum*. The false-positive test revealed all 35 markers as true sex-linked loci. Out of the 11 sex-linked SNP loci, we found one perfectly sex-linked in all individuals: all females are homozygous, suggesting XX biallelic form of the locus, while all males are heterozygous, suggesting an XY biallelic form for the locus. We found ten SNP loci that are moderately sex-linked, showing heterozygous (XY) allelic form in a few females, while homozygous (XX) allelic form in some males. However, none of the ten loci were discordant in more than 20% of individuals (as per the filtering criteria). Similarly, in PA loci, we found six perfectly sex-linked PA loci that are concordant (i.e., present in all males and absent in all females) in all 36 individuals (100%). We found slight discordance (<20%) in the rest of the 18 PA loci, showing the presence of these loci in a few females while absent in a few males (Table S1).

Given the number of individuals from each sex group (five males and three females) in cluster-2 was comparatively low, we identified markers that were 100% concordant to each sex as sex-linked. We found 38 SNPs and 51 PA loci that support a male heterogametic sex determination (XX/XY) system within this cluster (Figure 2). For instance, all SNPs were heterozygous in males and homozygous in females while all PA loci were present in males and absent in females (Table S2). In contrast, we found 54 SNPs and 67 PA loci that supports the female heterogametic sex-determination system (ZZ/ZW) (Figure 2): all the SNP loci were heterozygous and all the PA loci were present in females while they were all homozygous or absent in males (Table S2).

**Figure 2:**
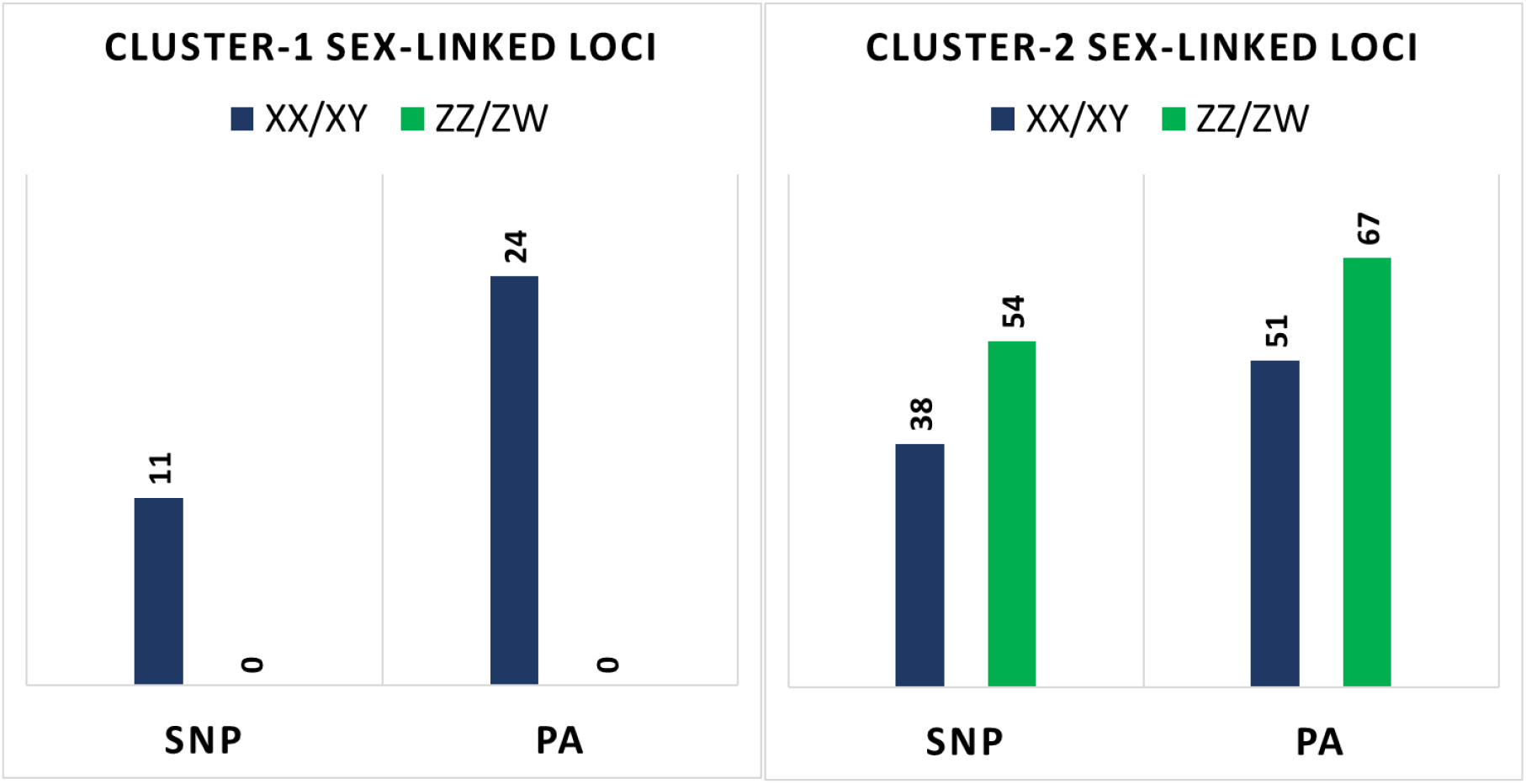
Number of sex-linked Single Nucleotide Polymorphism (SNP) and Presence-Absence (PA) loci in two clusters of *Platyplectrum ornatum*. XX/XY denotes markers supporting a male heterogametic sex determination system. ZZ/ZW denotes markers that support a female heterogametic sex determination system.

The pairwise Hamming distance matrix analysis using sex-linked SNPs and PA loci (Figure 3A-D) showed high differentiation between males and females in both clusters, suggesting a strong association of these markers to the putative sex chromosomes (Figure 3A-D). In cluster-1, the mean pairwise distance among males was 0.15 for the SNP loci and 0.11 for the PA loci. Among the female individuals of cluster-1, the mean pairwise distance was 0.05 for the SNP loci and 0.10 for the PA loci. The overall genetic distance between the two sexes was 0.97 for the SNP loci and 0.92 for the PA loci. Cochran–Armitage test verifies the significant association of all 11 SNP loci (χ2 = 0.97, p =0.04) and 24 PA loci (χ2 = 0.99, p =0.02) with phenotypic sex in cluster 1. In cluster 2, the mean pairwise distance among males was 0 for SNP loci and 0 for PA loci while the mean pairwise distance among females was 0 for SNP loci and 0 for PA loci. The overall genetic distance between the two sexes was 1 for the SNP loci and 1 for the PA loci. Cochran–Armitage test verifies the significant association of all 92 SNP loci (χ2 = 1, p =0) and 118 PA loci (χ2 = 1, p =0) with phenotypic sex in cluster-2.

**Figure 3:**
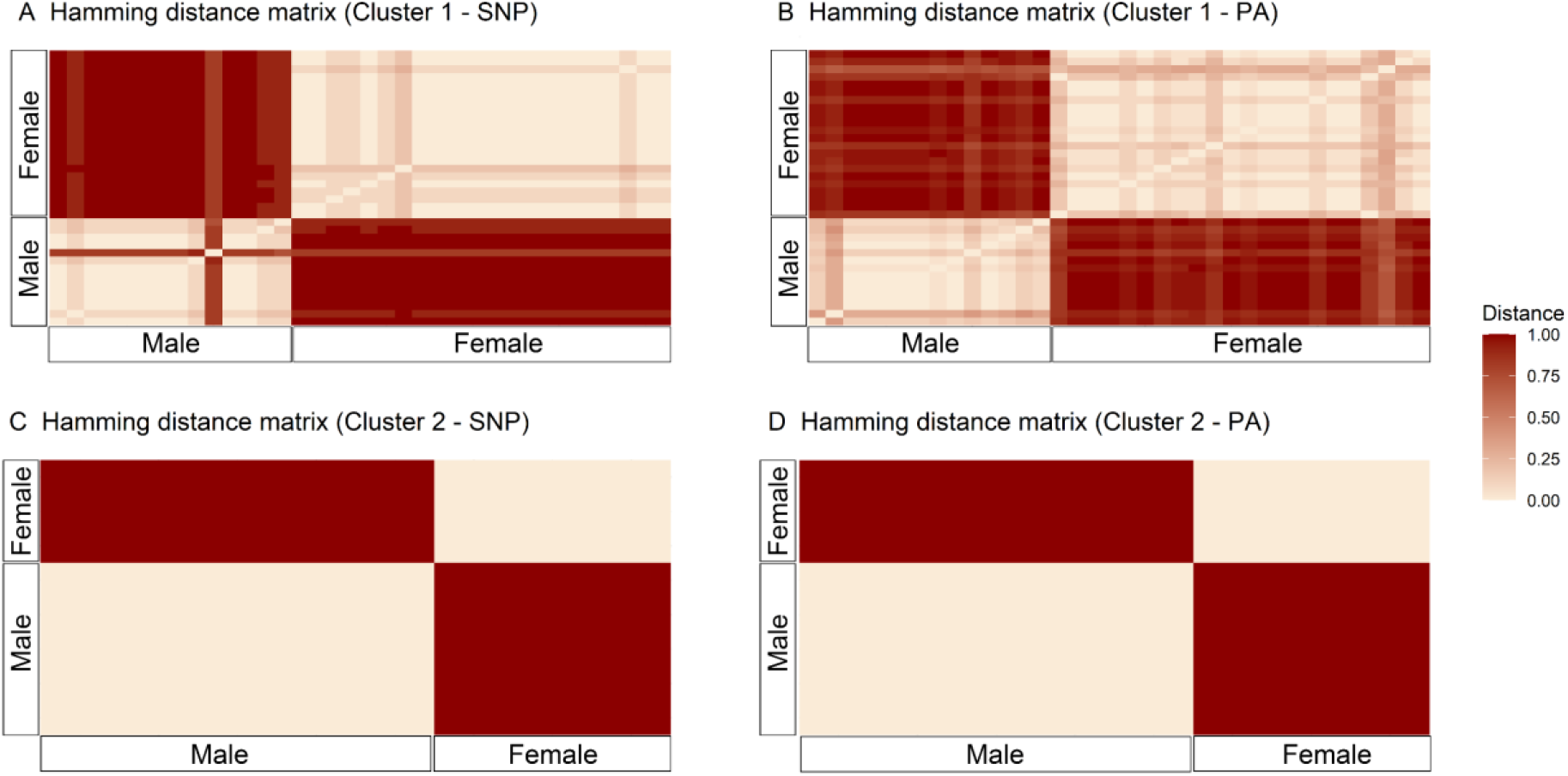
Hamming distance matrix using sex-linked SNP and PA loci in two clusters of *Platyplectrum ornatum*. A – genetic distance using SNP loci among individuals of cluster 1. B – genetic distance using PA loci among individuals of cluster 1. C – genetic distance using SNP loci among individuals of cluster 2. D – genetic distance using PA loci among individuals of cluster 2.

### Alignment of sex-linked markers against *P. ornatum* genome and BLAST search of the *P. ornatum* sex-linked scaffolds against *Rana temporaria* genome

We identified 29 sex-linked scaffolds in cluster-1 (Table S3) and 168 sex-linked scaffolds in cluster 2 (Table S4). The BLAST search of the sex-linked scaffolds against the *Rana temporaria* genome identified 7 genes in cluster-1 (Table S5) and 51 genes in cluster-2 (Table S6). Out of seven genes of the cluster-1, five genes (*PHLDA2, KCNC1, TBC1D17, Sacsin-like, PRMT1*) are located in chromosome (Chr) 10, one gene (*HS3ST6*) on Chr 6, and one gene (*MIPOL1*) on Chr 13 of *R. temporaria*. In cluster-2, we found a total of 7, 2, 6, 4, 5, 1, 3, 3, 12, 4, 1 and 1 gene harboured on Chr 1, Chr 2, Chr 3, Chr 5, Chr 6, Chr 7, Chr 8, Chr 9, Chr 10, Chr 11, Chr 12 and Chr 13 respectively (Table S6). Our analysis indicated that out of 51 genes, 23 genes were homologous to *P. ornatum* scaffolds that support a male heterogametic sex determination while 29 genes to that support a female heterogametic sex determination.

Findings from the BLAST search clearly indicate that the sex chromosome of *P. ornatum* is homologous to chromosome 10 of *R. temporaria*. Then, we further BLAST searched the sex-linked genes against the genome of *Xenopus tropicalis*. Our analysis found the 16 genes to be located on chromosome 7 of *X. tropicalis*. A comparative analysis between *R. temporaria* and *X. tropicalis* indicates that out of the 16 genes on chromosome 7 of *X. tropicalis*, 14 are located on chromosome 10 of *R. temporaria* while the other two are on chromosome 8 (Table S7).

### Cytogenetic screening of putative sex chromosomes in *Platyplectrum ornatum*

#### Karyotyping and C-banding

The karyotype analysis revealed 2n = 22 as described previously (Lamichhaney et al., 2021). Out of 11 homologous chromosome pairs, we found no heteromorphic sex chromosomes (Figure 4a-b) indicating *P. ornatum* has a homomorphic X and Y chromosome system (Figure 4a-b). Our C-banding analysis identified a comparatively higher accumulation of heterochromatin in the centromeric regions of all chromosomes along with a faint heterochromatin on either or both arms of chromosomes 1, 2, 3, 4, 5, and 6. We did not detect any sex specific C-banding pattern (Figure 4c-d).

**Figure 4:**
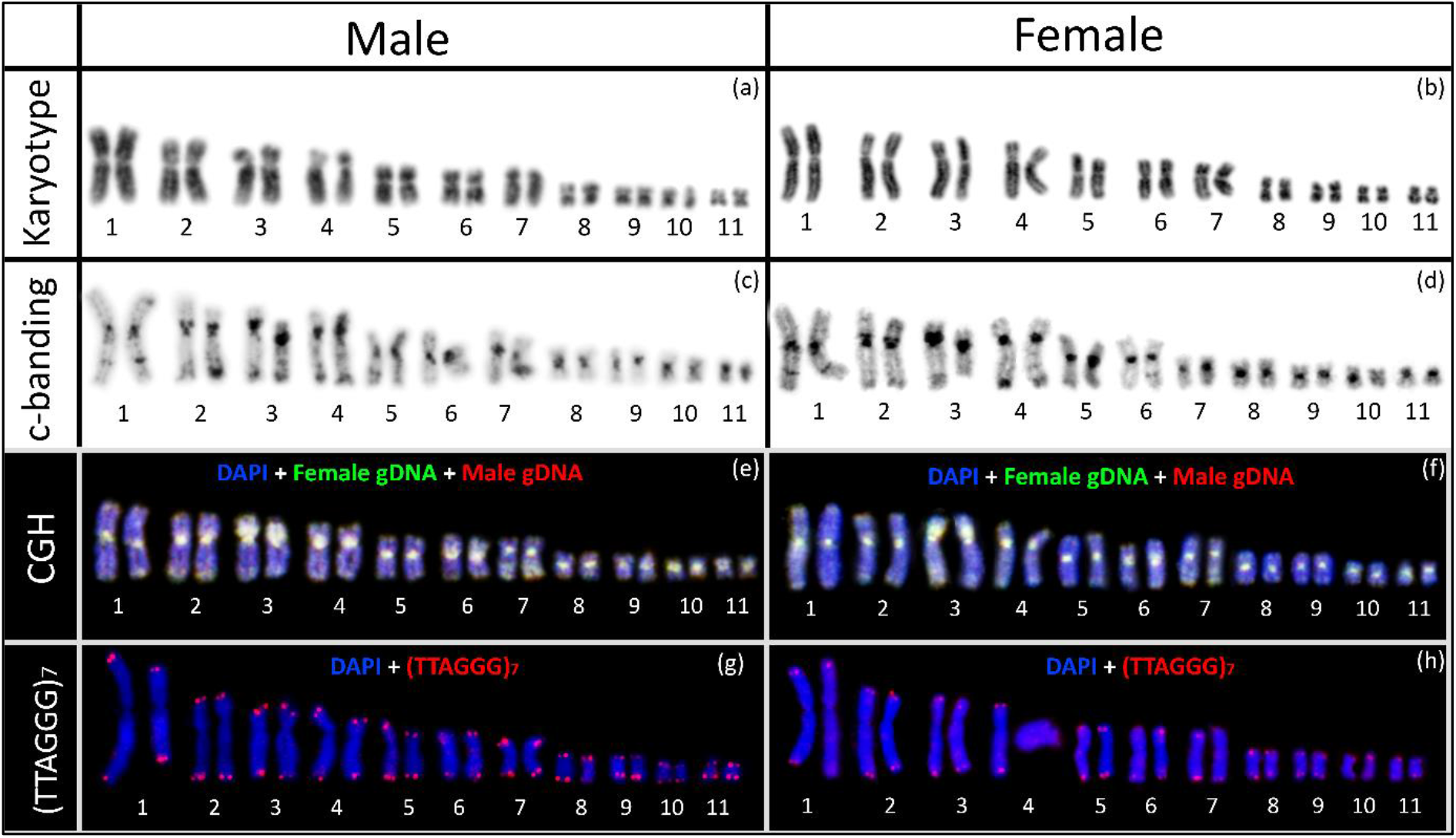
Cytogenetic screening of putative X and Y chromosomes in *Platyplectrum ornatum*. a) DAPI stained karyotype of male. b) DAPI stained karyotype of female. c) DAPI stained c-banding karyotype of male. d) DAPI stained c-banding karyotype of female. e) Comparative Genome Hybridisation (CGH) in male. f) Comparative Genome Hybridisation (CGH) in female. g) Fluorescence *in situ* Hybridisation (FISH) with telomere specific probes in male. h) Fluorescence *in situ* Hybridisation (FISH) with telomere specific probes in female.

#### Comparative Genome Hybridisation (CGH)

Our CGH analysis did not identify any sex specific pattern in *P. ornatum* (Figure 4e-f). We observed genome wide hybridisation pattern of fluorescently labelled DNA on both male and female chromosomes, indicating cytologically indistinguishable genomic differentiation between the X and Y chromosomes.

#### Fluorescence *in situ* Hybridisation (FISH) of Telomeric sequence

The FISH analysis indicates hybridisation of telomeric sequences only in the terminal region of each chromosome, but no sex specific interstitial telomere sequences (ITS) in *P. ornatum* (Figure 4g-h).

## DISCUSSION

### A new cryptic species of *P. ornatum?*

In this study, our aim was to identify the sex chromosomes and heterogametic sex in the Australian frog *P. ornatum*. Unexpectedly, however, we identified two genetically distinct clusters within the species based on SNPs. The F_st_ between the two is 0.24 and genetic distance (Nei) is 0.13, which are not high as a specific level, but PCA clearly indicates that they are genetically separated from each other and suggests establishment of reproductive isolation, probably, of pre-mating between the two, because they sympatrically inhabit the same region in Oyster cove. This finding provides an unexpected, well-suited model to study the early stage of sex chromosome evolution by comparison between the genetically closest two lineages and also contributes to understanding the mechanisms of speciation in Australian frogs.

### Homomorphic sex chromosomes and male or female sex determining mechanism in *P. ornatum*

As a new, cryptic species could be present in the two populations of *P. ornatum* studied here, we separated the two clusters-1 and -2 to individually isolate the sex-linked markers. From cluster-1, 35 sex-linked markers were isolated and were heterozygous in males (and homozygous in females), which are highly conserved markers across male individuals, suggesting that cluster-1 has male heterogametic sex determination. In contrast, cluster-2 had 210 isolated sex-linked markers, of which 89 were heterozygous or present in males (defined as male specific), while the other 121 were female heterozygous or present in females (female specific). The almost equal number of male and female specific markers indicates that cluster-2 has a homomorphic sex chromosome in both sexes with free recombination between the pair, and the heterogametic sex in cluster-2 remains to be solved. Through cytogenetic analyses of C-banding, CGH and telomere FISH, we demonstrated in cluster-1 of *P. ornatum* does not have any heteromorphic sex chromosomes and therefore, like many frogs, the two clusters of this species have homomorphic sex chromosomes with very little genomic differences between their sex chromosome pairs. This chromosome observation strongly supports the free recombination between the sex chromosomes: many moderately sex-linked markers in cluster-1 and equal numbers of male and female sex-specific markers in cluster-2.

The difference in number of sex-linked markers isolated between the two clusters may be due to either or both of the following two reasons. First, it is the depth of genetic variations across the individuals used for the analysis. The genetic distance within cluster-1 is 0.19 while it is 0.07 within cluster-2, suggesting much higher genetic homogeneity in cluster-2 even though they are collected from two distant locations. This homogeneity among the frogs may have made it easier to isolate many more sex-linked markers in cluster-2, similar to analysis using offspring of one-sibships. Second, the depth of genomic differentiation between the sex chromosome pair. In cluster-1, the genomic differences between the X and Y chromosome are at advanced stages of differentiation compared to that of cluster-2 X and Y and thus Y-linked markers are fixed. On the other hand, the recombination may more freely occur in the sex chromosome pair in cluster-2 except the sex-determining gene or region tightly linked to the sex determining locus.

### The sex chromosomes of *P. ornatum* are homologous to autosomal chromosome 10 of *Rana temporaria* and ZWY sex chromosome 7 of *Xenopus tropicalis*

Using the recently published *P. ornatum* genome (Lamichhaney et al., 2021), we assigned 219 out of 245 sex-linked markers to *P. ornatum* scaffolds (Table S3 and Table S4). Compared to the DArTseq loci, these scaffolds are larger in size (bp) and allowed us to identify sex chromosome homologies across multiple frog species with chromosome level assembly. We identified 7 genes from cluster-1 and 51 genes from cluster-2 that are included in the sex-linked scaffolds. The comparative analysis across two frog species with well annotated chromosome level genome assembly indicates that the 14 genes mentioned above are located in chromosome 10 of *Rana temporaria* and those plus two more genes are in the sex chromosome of *Xenopus tropicalis* (Chromosome 7) (Fig 5). The sex chromosome 7 of *X. tropicalis* is partially homologous to the ZZ-ZW type of sex chromosome 7 in the Japanese bell ring frog *Buergeria buergeri* (Uno et al., 2015). The two chromosomes share two genes, *Cyp17* and *Got1*, which are also mapped on chromosome 9 of the Japanese soil-frog *Glandirana rugosa* (Sakurai et al., 2008; Suda, Uno, Mori, Matsuda, & Nakamura, 2011), and one of which (*Cyp17*) is on chromosome 8 of *Rana temporaria*. In addition, the late replication banding patterns are highly conserved in the chromosomes 9 as well as the other chromosomes with no apparent chromosomal rearrangements between the latter two species (Miura, 1995). Therefore, these four chromosomes of four species are partially or wholly homologous to each other. In contrast, the 14 genes identified in the sex chromosomes of *P. ornatum* are located on chromosome 10 of *R. temporaria*, but not on chromosome 8 that is homologous to a part of chromosome 7 of *X. tropicalis* (Fig. 5). This could be explained by the chromosomal arrangements that specifically occurred in *X. tropicalis*, of which chromosome 7 was derived from a fusion of two chromosomes, one of which is homologous to chromosome 8 and the other to chromosome 10 of *Rana temporaria* (Fig.5). Of particular interest is that the sex determining region of *X. tropicalis* is estimated to be located within 10.4 Mb from the terminal tip of the sex chromosome (Furman et al., 2020). The two genes of *Cyp17* and *Got1*, which are located within the non-recombining region in sex chromosome 7 of *B. buergerii*, are located next to the sex determining region on Z, W and Y chromosomes. The two sex-linked genes, *GAPDH like* and *DPYSL4* of *P. ornatum*, are located at or around the sex determining region of chromosome 7 of *X. tropicalis* (Fig. 5). Thus, it is likely that the sex determining gene of *P. ornatum* is shared with the sex determining gene(s) of *X. tropicalis* and *B. buergerii*. Probably, cluster-1 of *P. ornatum* has male heterogametic sex determination, whereas we have not enough data to decide the heterogametic sex in cluster-2. Interestingly, chromosome 7 of *X. tropicalis* can work as a W chromosome to determine femaleness or Y chromosome to determine maleness (Furman et al., 2020; Roco et al., 2015). Therefore, the sex-determining locus in the sex chromosome of cluster-2 of *P. ornatum* has a potential to dominantly determine male and/or female.

**Figure 5.**
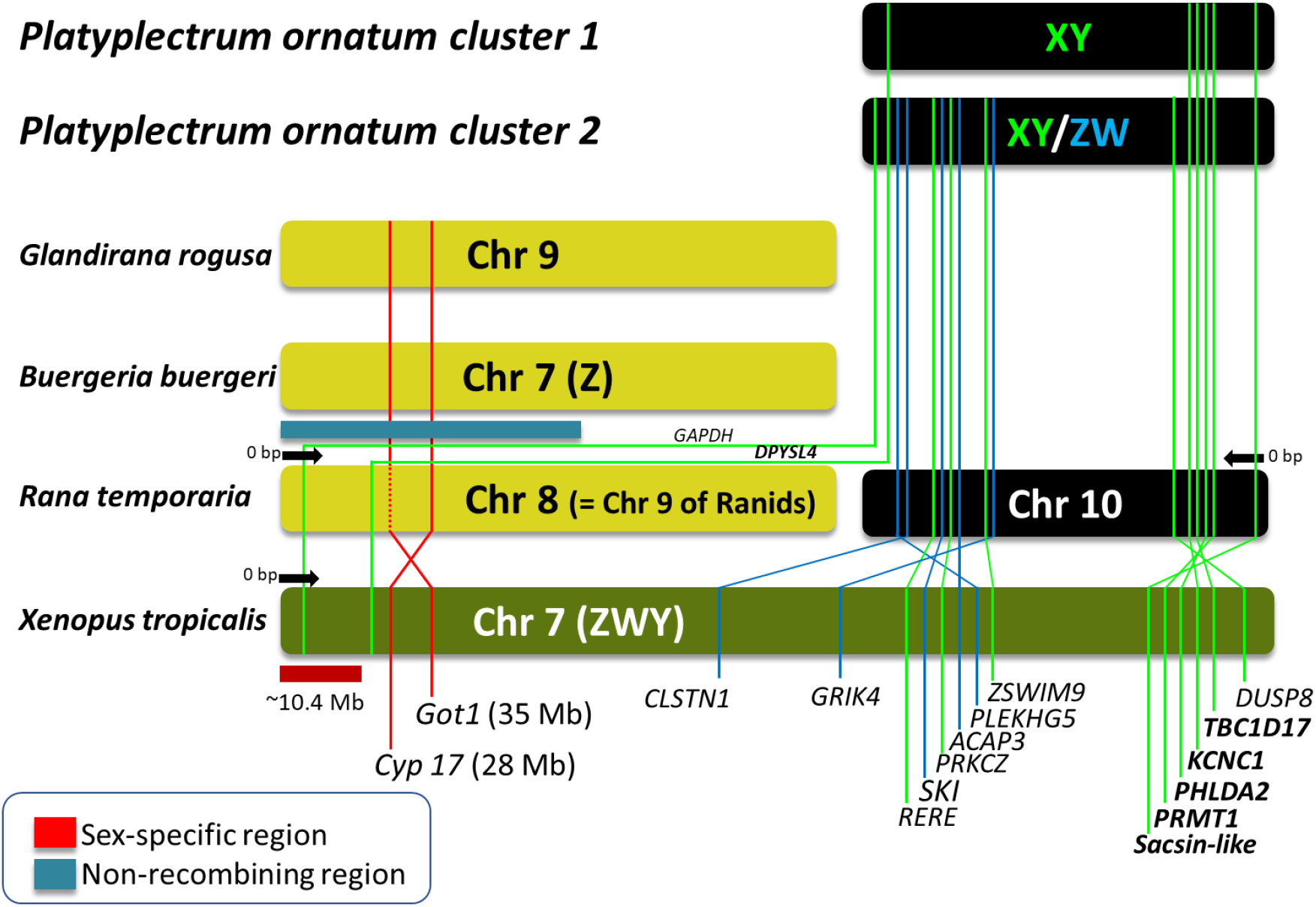
Diagram showing sex chromosome homology of *P. ornatum* with chromosomes of other frog species. The sex determining region (10.4 Mb) of *X. tropicalis* is estimated to lie at the terminal tip of sex chromosome 7 (Furman et al. 2020), indicated by a red bar, and two genes (*Cyp17* and *Got1*) are located next to it. Likewise, chromosome 8, 9 and ZW sex chromosome 7 of *Glandirana rugosa, Rana temporaria* and *Buergeria buergeri*, respectively, share the two or one genes, and are located within the non-recombining region in B. buerugeri, indicated by a dark blue bar. The six and 14 sex-linked genes identified in cluster-1 and -2 of *P. ornatum*, respectively, are shared with chromosome 10 (two with chromosome 8) of *Rana temporaria* or sex chromosome 7 of *X. tropicalis*.

### Sex chromosome evolution in Australian frogs

In Australia, out of 248 native frog species described, 133 species have had their karyotypes examined to date (King, 1980, 1990). Surprisingly, a heteromorphic sex chromosome pair has been identified in just one species, *Crinia bilingua* (Mahony, 1991); the smallest chromosome pair out of 12 haploid complements is a ZZ-ZW type of heteromorphic sex chromosomes. Focussing on the genetic, ecological and geographic characteristics of this species and comparing it to other Australian frog lineages might better help us to understand what drives the evolution of sex chromosomes from homomorphy to heteromorphy. One of the most intriguing questions that remains is how often the sex chromosome turnover has been repeated (or, inversely have not), during the phylogenetic history among Australian native frog lineages, which explosively and rapidly adapted to Australia’s unique climate and habitats and conserved almost completely homomorphic sex chromosomes. Thus, future studies using such DArT molecular techniques as we have outlined here are expected to find new and unique cases in sex chromosome evolution and will contribute to understanding the mechanisms underpinning this.

## MATERIALS AND METHODS

### Ethics statement

All collection and animal handling were performed following the approved animal ethics by NSW scientific permit SL101269, Macquarie University animal ethics approval 2019-010 and University of Canberra animal ethics committee (AEC 14-09 and AEC 18-01). Tissues from all 44 individuals were stored at -20°C until further processing.

### Animals and phenotypic sexing

We used 44 phenotypically sexed individuals of *Platyplectrum ornatum* for genotyping by sequencing (GBS). Nineteen (19) phenotypic males and 25 phenotypic females were collected between January 2016 to October 2018 from two sites: Oyster Cove and Dudley, NSW, Australia (Figure 1). Phenotypic sex was determined in the field by a combination of the presence or absence of male nuptial pads, presence or absence of an enlarged flange on the second digit of the hand used by females to beat egg masses into a frothy mass and by throat colouration (Clulow & Swan, 2018). We further confirmed phenotypic sex in the lab by dissection and inspection of the gonads post-euthanasia.

### Genotyping by sequencing using Diversity Arrays Technology Sequencing (DArTseq)

Approximately 25 mm of muscle tissue was collected from each individual and submitted to Diversity Arrays Technology (DArT) Pty Ltd (University of Canberra, Bruce, ACT, Australia) for genotyping by sequencing (Kilian et al., 2012). The genomic DNA was extracted by DArTseq™ following proprietary manufacturer’s instruction. Genomic DNA quality was confirmed by running a 1.2% agarose gel electrophoresis. DArTseq™ uses a combination of complexity reduction methods and next-generation sequencing (NGS) to generate thousands of Single Nucleotide Polymorphisms (SNP) and SilcoDArT (presence-absence) loci. Approximately, 100 ng of DNA from each sample was digested using *Pst*I and *Sph*I restriction enzyme. Digested DNA were subjected to a ligation reaction using a *PstI* compatible adaptor (consisting of an Illumina flow cell attachment sequence, sequencing primer sequence and a unique barcode sequence) and a *SphI* compatible adaptor (consisting of an Illumina flow-cell attachment region). The ligated fragments then underwent 30 rounds of PCR (94°C for 20 seconds, 58°C for 30 seconds and 74°C for 45 seconds), followed by an extension of seven minutes at 72°C. Following PCR, equimolar amounts of amplification products derived from each individual were bulked and applied to Illumina’s proprietary cBlot bridge PCR, which was followed by sequencing on an Illumina Hiseq2000. The single read sequencing was run for 77 cycles.

The raw sequences generated by Illumina Hiseq2000 were further filtered based on reproducibility average values, the read depth of each sequence, Polymorphism Information Content (PIC) and call ratio of each sequence across all individuals. The final SNP and PA data were then converted into a csv file containing 18 loci matrices including AlleleID, CloneID, AlleleSequence, TrimmedSequence (sequence after removing adapters), SNP (the polymorphic neucleotide)SnpPosition, CallRate (proportion of individual for called for a particular locus), OneRatioRef (presence of the reference allele), OneRatioSnp (presence of alternate allele), FreqHomRef (proportion of individuals with homozygous to reference allele), FreqHomSnp (proportion of individuals with homozygous to SNP allele), FreqHets (proportion of individuals with presence of both alleles), PICRef (polymorphic information content of the reference allele), PICSnp (polymorphic information content of the alternate allele), AvgPIC (average PIC), AvgCountRef (average count of the reference allele), AvgCountSnp (average count of the alternate allele) and RepAvg (reproducibility average of the locus). Each locus presented as “0” alternatively known as “homozygous reference”, “1” alternatively known as “homozygous SNP” and “2” alternatively known as “heterozygous”. If an allele failed to call successfully for an individual due to sequencing error or low-quality genomic DNA, the locus was presented as a null allele or “-”. For PA markers, the presence of a marker was presented as “1” and the absence of the marker in the genomic representation was referred as “0”. Similar to the SNP loci, null alleles for PA markers were also presented as “-”.

### Population genetic structure analysis

We performed population genetic structure analysis to identify genetic variation among all 44 individuals analysed in this study. We filtered out all sex-linked loci and loci that were not with 100% call ratio and 100% reproducibility average for this analysis. The filtering of loci was performed using “dartR” version 1.9.4 package in R (Gruber, Unmack, Berry, & Georges, 2018). The Principal Component Analysis (PCA) was performed using “dartR” version 1.9.4 package in R (Gruber et al., 2018). The genetic structure analysis was performed with an F_st_ based structure analysis software “Structure” version 2.3.4. We performed an admixture model with 1000 burn in and 1000 MCMC assuming 10 populations. Each iteration was run 10 times to identify appropriate K value among the data set. The deltaK was calculated using “Structure Harvester” (Earl, 2012).

### Identification of sex-linked markers

We combined previously published pipelines to identify both single nucleotide polymorphism (SNPs) and presence-absence (PA) sex-linked loci in this study (Hill, Burridge, Ezaz, & Wapstra, 2018; Jeffries et al., 2018; Lambert, Skelly, & Ezaz, 2016; Nguyen et al., 2021; Sopniewski et al., 2019). First, we filtered out all loci (both SNPs and PA) that were below 80% call ratio. The remaining loci were tested for the presence of both male heterogametic (XX/XY) and female heterogametic (ZZ/ZW) sex determination systems. For SNPs, a marker was identified as sex-linked if it was at least 80% heterozygous in one sex and at least 80% homozygous in the opposite sex. For PA loci, we applied the same filtering criteria as SNPs where markers that were present in a minimum of 80% in one sex and absent in minimum of 80% in the opposite sex are considered as sex-linked. However, this pipeline tends to identify a small number of SNPs that support the filtering criteria but are not true sex-linked loci particularly if the representative samples from each sex is low, i.e., below 13 individuals per sex as suggested by Lambert et al. (2016). To filter out such loci, we performed a false positive test across all identified sex-linked loci by calculating the proportion of homozygous alleles as described by Jeffries et al. (2018). For example, in a male heterogametic sex determination system (XX/XY), all true sex-linked loci will show a minimum of 80% homozygosity (either reference or alternate alleles for SNPs) to female or the homogametic sex. Similarly, in a ZZ/ZW system, all true sex-linked loci will show a minimum of 80% homozygosity (either reference or alternate) to male or the homogametic sex.

We used the “countif” function in Microsoft Excel for identification of sex-linked markers. For calculating Pairwise genetic distance (Hamming distance matrix) we used the “rdist” function in the “rdist” package in “R” version 3.6.2 (R Core Team, 2017). The Cochran– Armitage trend test (CATT) was performed using R package “CATT” version 2.0 (R Core Team, 2017).

### Alignment of sex-linked markers against *P. ornatum* genome and orthology analysis using BLAST search

DArTseq loci are usually a result of random sampling from the genomic representation and comparatively small sized (69 bp or less). Hence, a priori knowledge about gene association with individual alleles are unknown (Shams et al., 2019). To identify homologous genes and sequences, we mapped all sex-linked SNP and PA loci to the recently published *P. ornatum* genome assembly (Lamichhaney et al., 2021) and used the *P. ornatum* sex-linked scaffolds for a Basic Local Alignment Search Tool (BLAST) analysis. The mapping of sex-linked SNP and PA loci to the *P. ornatum* genome was performed with a Medium/Fast sensitivity using “Geneious” version 10.2.6 (Kearse et al., 2012). The homologous scaffolds were subject to BLAST searches against the well annotated (chromosome level) frog genomes for *Rana temporaria* and *Xenopus tropicalis*. We used NCBI BLASTn tools (http://ncbi.nlm.nih.gov/Blast.cgi) to perform a megablast search with a threshold e-value 10^−3^.

### Molecular Cytogenetics analysis to identify sex chromosome pairs

#### Metaphase chromosome preparation

To identify putative sex chromosomes in *P. ornatum* we performed molecular cytogenetic analyses in one male and one female. Metaphase chromosomes were prepared following the protocol described in (Netto, Pauls, & de Mello Affonso, 2007) with slight modification. Briefly, bone marrow was extracted and rinsed in a small glass Petri dish with 5 ml chilled Roswell Park Memorial Institute (RMPI) 1640 (Gibco, chilled at 4°C) culture medium and then transferred to a centrifuge tube containing 10 ml RPMI. Approximately three drops of colchicine (0.1%w/v) was then added to the solution and incubated for 45 minutes at room temperature. The cell suspension was then centrifuged at 1000 RPM for 10 minutes. Supernatant was discarded and 10 ml hypotonic solution (0.075 M KCl) was added, mixed and incubated the cell suspension for 40 minutes at room temperature. Five drops of chilled freshly prepared Carnoy’s solution (methanol:acetic acid 3:1 at 4°C) was then added to the cell suspension and was centrifuged at 1000 RPM for 10 minutes. After centrifugation supernatant was discarded and cells were fixed in freshly prepared Carnoy’s solution at room temperature. Fixed cells were centrifuged three times at 1000 RPM for 10 minutes, discarding the supernatant after each centrifugation step. After the last centrifugation, supernatant was discarded and cells pellet was resuspended with Carnoy’s solution at a ratio of 1:1 (v/v) and mixed the solution to achieve a homogeneous cell suspension. Cell suspensions were dropped onto glass slides and air-dried. For DAPI (40-6-diamidino-2-phenylindole) staining, slides were mounted with antifade medium Vectashield (Vector Laboratories, Burlingame, CA, USA) containing 1.5 mg/mL DAPI.

#### C-Banding

Detection of heterochromatin (C-banding) is a common technique in identifying sex chromosomes. We performed C-banding on metaphase chromosomes of both the male and female to identify sex specific heterochromatinisation following the protocol described in Ezaz et al. (2005) with a slight modification. Briefly, 20–25 µL of cell suspension was dropped on slides, air dried and aged at -80°C overnight. Slides were then treated with 0.2 M HCl at room temperature for 20 minutes followed by 5% Ba(OH)_2_ at 45 °C for 3 minutes. The slides were then incubated in 2× saline sodium citrate (SSC) at 65 °C for 60 min. Finally, the chromosomes were stained with antifade medium Vectashield containing 1.5 mg/mL DAPI.

#### Comparative Genomic Hybridisation (CGH)

Hybridisation of fluorescently labelled genomic DNA from each sex is one of the most effective and widely used techniques to identify sex chromosome pairs in a species. We performed Comparative Genomic Hybridisation (CGH) on both male and female metaphase following the protocol described by Ezaz et al. (2005). Briefly, female and male total genomic DNA was labelled by Nick translation (NT), incorporating Spectrum-Green and Spectrum-Red dUTP, respectively, NT kit (Abbott Molecular, Macquarie Park, Australia). Nick translation reaction was incubated at 15°C for 2 hours. After the incubation, NT labelled DNA were checked using 1% agarose gel electrophoresis for size fractionation to 200–500bp. Labelled DNA were co-precipitated by overnight incubation at −20°C with 5–10 µg of salmon sperm DNA, 20 µg glycogen and 3 volumes of chilled 100% ethanol. Precipitation reaction was centrifuged at maximum speed for 30 min, and the supernatant was discarded and air-dried. Depending on the size of pellet, the co-precipitated probe DNA was resuspended in 30–40 µL hybridisation buffer (50% formamide, 10% dextran sulfate, 2× SSC, 40 mmol/L sodium phosphate pH 7.0 and 1× Denhardt’s solution). The hybridisation and labelled DNA solution was added to the glass slide containing aged metaphase chromosomes, covered with a cover slip and sealed with rubber cement. Labelled DNA and metaphase chromosomes were denatured at 70°C for 5 minutes and incubated at 37°C for 3 days in a humid chamber. Slides were washed once at 60°C in 0.4× SSC, 0.3% Igepal for 3 minutes, followed by another wash at room temperature in 2× SSC, 0.1% Igepal for 2 minutes. Slides were then air-dried and mounted with antifade medium Vectashield containing 1.5 mg/mL DAPI. Images were captured using a Zeiss Axioplan epifluorescence microscope equipped with a charge-coupled device (CCD) camera (RT-Spot), (Zeiss, Oberkochen, Germany) using filters 02, 10 and 15 from the Zeiss fluorescence filter set or the Pinkel filter set (Chroma technologies, filter set 8300, Bellows Falls, VT, USA). ISIS scientific imaging software (Metasystems, Altlussheim, Germany) was used for image capture and analysis, including karyotyping. For FISH and CGH image analysis, multiple functions of ISIS scientific imaging software were used, such as signal normalizing and background correction. Processed images were pseudocoloured and superimposed, using ISIS scientific imaging software.

#### Fluorescence *in situ* Hybridisation using telomeric sequences

To identify interstitial telomeric sequences, we performed Fluorescence *in situ* Hybridisation (FISH) using vertebrate specific telomeric repeats. The ready to use 5’-Cy3-labeled single strand telomeric probes (TTAGGG)_7_ (PNA Bio) were purchased from PNAGENE Inc (Daejeon, Korea). The FISH was performed following the manufacturer protocol. Briefly, air dried slides with metaphase chromosomes were dehydrated in a cold ethanol series (70% (v/v), 85% (v/v) and 100% (v/v) ethanol) for 1 minute each. Air dried slides were preheated at 80°C for 5 minutes. For each slide, 20 µl (150-200 ng) probe and hybridisation buffer mixture was added and sealed with a cover slip. Chromosomes were then denatured at 85°C for 10 minutes and incubated in a dark chamber at room temperature for 60 minutes. After 60 minutes, the cover slips were removed, and the slides were washed in a solution containing 2×SSC and 0.1% (v/v) Tween-20 at 60°C for 10 minutes. This step was performed twice before washing slides again at room temperature for 1 minute. Slides were then air dried and mounted with antifade medium Vectashield containing 1.5 mg/mL DAPI.

## SUPPLEMENTARY DATA

Supplementary data to this article can be found online.

## AUTHOR CONTRIBUTIONS

TE conceived the idea. SC performed field work and TE sample collection. CS, FS and ZM conducted lab works. CS, FS, IM and TE performed the analysis. CS, FS, TE and IM wrote the initial draft. All co-authors contributed intellectually to writing and editing the draft multiple times. All authors read and approved the final version of the manuscript.

## ACKNOWLDGEMENTS

The authors would like to acknowledge Julie Strand for assistance with tissue sub sampling.

